# Disentangling bidirectional relationships between glucocorticoids and behavior: Experimentally elevated corticosterone levels correlate with rapid, sex-specific changes in food-acquisition behaviors of food-limited seabird chicks

**DOI:** 10.1101/2025.07.14.664366

**Authors:** Z Morgan Benowitz-Fredericks, Alexis P. Will, Sierra N. Pete, Stephanie Walsh, Shannon Whelan, Alexander S. Kitaysky

## Abstract

Bidirectional relationships between hormones and behaviors are hypothesized: i.e. energetic consequences of behaviors may alter glucocorticoid levels, and elevated glucocorticoids may alter behavioral responses to stimuli in context-appropriate ways. We tested these relationships between the avian glucocorticoid corticosterone and behaviors related to food-acquisition in free-living seabird chicks (Black-legged kittiwakes, *Rissa tridactyla*) with robust HPA-axis activity. First, we tested the hypothesis that behavior affects HPA axis activity by quantifying chick behavior for 60 min, then taking a blood sample at baseline and 15-min post-restraint. Only feeding predicted subsequent corticosterone levels (fewer feeds = higher restraint-induced corticosterone). However, the hypothesis that increased corticosterone secretion alters behavior was supported: post-restraint corticosterone levels were positively correlated with increases in begging and aggression in the subsequent hour, primarily in males. In a second experiment, we isolated the role of corticosterone from other components of the stress response by administering exogenous topical corticosterone, which confirmed a positive relationship between subsequent corticosterone levels and changes in aggression, but not begging or feeding. Corticosterone:behavior relationships depended on nutritional context - they were eliminated in nests with experimental food supplementation. Finally, higher levels of corticosterone in response to restraint were associated with more rapid elimination of siblings, presumably increasing chick direct fitness by eliminating competition. We conclude that the magnitude of the endogenous corticosterone response to challenges is a) negatively correlated with recent feeding rates, b) subsequently associated with rapid, sex- and context-specific changes in chick behaviors, at least some of which are likely causally- driven by elevated corticosterone, and c) associated with fitness-relevant consequences.

## 1. INTRODUCTION

Hormones can change behavior rates by altering the probability that an animal will respond to a particular context and set of stimuli with appropriate behaviors, while behavior itself can also alter hormone levels (Adkins-Regan, 2005; Hau and Goymann, 2015; Myers et al., 2014, Wingfield et al. 1990). These bidirectional relationships have been well-established in the context of competitive or reproductive behaviors and androgens in vertebrates (Casto and Edwards, 2016; George and Rosvall, 2022; Goymann, 2009; Olivera, 2004; Pradhan et al., 2015, Rosvall et al., 2020), where they often differ by sex (Jennings and DeLecia, 2020; Smiley et al., 2022). However, glucocorticoid hormone levels can also influence and be influenced by behavior (Haller, 2014; Horton and Hollberton, 2009; Myers et al., 2014). Glucocorticoids’ effects are usually framed in terms of context-appropriate redirection of energy towards behaviors that will help an animal survive a threat and re-establish homeostasis (Orchinik, 1998, Sapolsky, 2000, Wingfield and Kitaysky, 2002). Simultaneously, behaviors require energy expenditure and may thereby affect the activity of the hypothalamo-pituitary-adrenal (HPA) axis, the endocrine cascade culminating in glucocorticoid release (Falson et al., 2009; Grunst et al., 2024; Hackney and Waltz, 2013; Mateos, 2005). Tests of these bidirectional glucocorticoid:behavior relationships in free-living animals are rare, and when relationships exist, they may be modified by a number of factors that warrant investigation.

In birds, sex, food availability and time scale may all modulate relationships between corticosterone (the primary avian glucocorticoid) and behavior. First, relationships between behavior and corticosterone may differ by genetic sex, because animals may have sex-specific strategies for coping with challenges (Ouyang et al., 2012; Wingfield, 1985). Second, the relationships between physiology and behavior may be altered by the environmental context (Killen et al., 2013; Pradhan et al., 2015), particularly food availability. Food availability has consistently been shown to affect both behavior (Golabek et al., 2012; Quillfelt and Masello, 2004;) and HPA activity (reviewed in Busch and Hayward, 2009; Kitaysky et al., 2007; Sorensen et al., 2017) in birds. The behavioral and metabolic responses to elevated glucocorticoids may differ in individuals depending on constraints imposed by their energy budgets. Thus, by altering the amount of resources available to allocate across competing demands, variation in food availability may either obscure or reveal underlying relationships between physiology and behavior (Benowitz-Fredericks et al., 2022; Killen, et al. 2013). Finally, temporal scale is likely to be important. Despite animals’ capacity to increase glucocorticoids on the scale of minutes (Romero and Reed, 2005), most studies that measure or manipulate glucocorticoids evaluate its relationship with behavior on the scale of days, potentially missing an important shorter-term time-window when glucocorticoids and behavior may interact. In this study, we assessed the effects of food availability and genetic sex on glucocorticoid:behavior relationships over a short time scale in young black-legged kittiwakes (*Rissa tridactyla),* a species whose chicks have robust adrenocortical activity and intense sibling competition.

### Black-legged kittiwakes

We conducted two experiments to test hypotheses about relationships between corticosterone and a set of focal behaviors related to food acquisition (begging, aggression and feeding) on a short time scale (an hour). We studied five-day old nest-bound chicks that engage in intense sibling competition and have adult-like hypothalamo-pituitary-adrenal (HPA) axis activity (Benowitz-Fredericks et al., 2024). Kittiwakes are long-lived seabirds with broods of 1-3 chicks who engage in facultative siblicide, which peaks in the first 10 days post-hatch (White et al., 2010). When siblicide occurs, the older “A” chick eliminates its younger siblings (“B” or “C” chick) by physically attacking them. The attacks may prevent the younger sibling from acquiring food, drive them from the nest or kill them outright (Dickins, 2021, Braun and Hunt, 1983, White et al., 2010). While it can be difficult to confidently ascertain the cause of death for kittiwake chicks, which often simply disappear from the nest between twice-daily nest checks (*pers. obs.*), rates of aggression and brood reduction both increase when food availability is low (Vincenzi et al., 2015; White et al., 2010; Gill et al., 2002; Suryan et al., 2002), suggesting that low food availability increases siblicide. Low food availability also leads to increased secretion of corticosterone in kittiwake chicks (Kitaysky et al., 1999; Benowitz-Fredericks et al., 2024), suggesting that corticosterone levels may affect the probability of engaging in behaviors associated with food acquisition and/or siblicide. By 5 days post-hatch, kittiwake chicks engage in intensive begging and aggression (White et al. 2010), have sexually dimorphic developmental trajectories (Merkling et al., 2012) that may be associated with sex differences in hormone:behavior relationships, and have rapidly responsive HPA axes that are sensitive to food availability (Benowitz-Fredericks et al., 2024). These traits make them good candidates for testing hypotheses about sex-specific relationships among corticosterone, behavior and food availability on short time-scales.

### Study hypotheses

In this study, we investigate the bidirectional relationship between behavior and corticosterone in black-legged kittiwake chicks, which may shed light on why such young, nest-bound chicks are capable of rapidly and dramatically elevating corticosterone. This question is important because many studies have indicated that there are costs to chicks of exposure to elevated glucocorticoids during development, presumed to reflect unavoidable trade-offs (Herborn et al., 2014; Kitaysky et al., 2003; reviewed et Schoech et al., 2011), while benefits are often assumed but rarely tested, especially in free-living animals. No clear pattern has been identified to explain why the HPA-axes of some species are hyporesponsive early in development while others can mount robust responses at very young ages (Benowitz-Fredericks et al., 2024), but behavioral demands may play a role. Increases in begging and aggression have been identified as potential benefits of elevated corticosterone in chicks (Kitaysky et al., 2001; Kitaysky et al., 2003; Wada et al. 2008). While this is consistent with our general understanding of glucocorticoids as hormones that, during challenges, redirect allocation of energy towards physiology and behaviors in ways that maximize fitness (in this case, behaviors associated with increasing food acquisition), most studies have evaluated effects of chronic or repeated exposure to elevated corticosterone on the scale of days to weeks, rather than short-term effects, despite the rapid dynamics of HPA axis activity. In addition, few studies have considered chick sex, and none have manipulated food availability for free-living birds.

Using two different experiments, we examined potential causes and correlates of HPA-axis activity in young kittiwake chicks. We tested a set of four broad, non-exclusive hypotheses in male and female kittiwake chicks. 1) Because engaging in behavior expends energy, and corticosterone levels can reflect metabolic activity (Jimeno and Verhulst, 2023), we tested the proximate hypothesis that recent chick behavior drives HPA axis activity. 2) Because glucocorticoids are assumed to redirect allocation of energy towards context-appropriate behaviors, we tested the hypothesis that HPA axis activity promotes rapid changes in behavior in ways that facilitate food acquisition. 3) Because food availability can alter relationships between hormones and behavior (Cote et al., 2010, Killen et al., 2013, Benowitz-Fredericks et al., 2022), and corticosterone in kittiwake chicks rises when food availability is low (Kitaysky et al., 1999, Kitaysky et al., 2001, Benowitz-Fredericks et al., 2024), we also tested the hypothesis that food availability modulates hormone:behavior relationships. 4) Finally, we tested the ultimate hypothesis that young kittiwake chicks have highly reactive HPA axes because HPA axis activity is relevant to fitness (as measured by rapid elimination of competition from a sibling - the occurrence and speed of brood reduction).

### Focal behaviors

We focused on behaviors that we considered relevant to food acquisition (and therefore glucocorticoid secretion) in young chicks: begging (a nest-bound chick’s primary mode of ‘foraging’ by soliciting food from parents), aggression (acquiring a larger share of food by suppressing or eliminating competition from younger siblings), and feeding (an integration of both chick and parent behaviors). Most studies of corticosterone and chick behavior have focused on begging, but chicks of siblicidal species like kittiwakes have larger behavioral repertoires than those who compete only through begging, and behaviors like aggression may also interact with HPA axis activity on acute time scales (Kitaysky et al., 2003).

### Elevating endogenous and exogenous corticosterone levels in seabird chicks

The first experiment addressed hypotheses 1 - 4 by examining correlative relationships between behavior and endogenous corticosterone levels (baseline and following 15 min of restraint) over the period of two hours.

To test Hypothesis 1, we evaluated the relationships between behavior over the course of an hour and subsequent baseline corticosterone levels, measured at the end of that hour. If the hypothesis that recent behavior drives HPA axis activity in young chicks is true, we expect relationships between aggression, begging, and/or feeding and corticosterone levels at the end of the hour of behavior. To test Hypothesis 2, that elevated corticosterone rapidly alters behavior, we evoked endogenous elevations in corticosterone via 15 min of handling and restraint, and tested relationships between corticosterone levels (baseline and post-restraint), and changes in behavior in the subsequent hour. The prediction of this hypothesis is that the magnitude of circulating corticosterone levels should be associated with the magnitude of changes in behavior.

We tested Hypothesis 3, that relationships between corticosterone and behavior are stronger when food availability is limited, by comparing these hormone:behavior relationships in both control and experimentally food-supplemented nests. Finally, we tested Hypothesis 4 with an indirect measure of fitness: the survival of the younger chick. If HPA axis activity is related to fitness, we would expect that chicks with higher corticosterone levels would eliminate their competitors (younger siblings) faster.

In the second experiment we addressed the possibility that correlative relationships between behavior and elevated endogenous corticosterone levels evoked by restraint (as predicted by Hypothesis 2, see above) may be driven by components of the endogenous stress response other than corticosterone. We experimentally decoupled the endogenous stress response from corticosterone elevation by quantifying behavior in chicks from non-food-supplemented nests for one hour before and one hour after experimentally elevating exogenous corticosterone using minimally-disruptive topical hormone application. Comparison to a sham-treated control group allows us to attribute any treatment differences to the effects of corticosterone alone. To the best of our knowledge, this is the first study to test short-term hormone:behavior relationships using both endogenous and exogenous elevations of corticosterone in free-living birds.

## METHODS

### 2.1 Field site and focal chicks

All research was conducted at a colony on Middleton Island, AK (59.433552, -146.341008), where free-living kittiwakes nest on the face of a modified radar tower. Each nest site is equipped with a sliding, one-way mirrored glass window that allows nest observation and access from inside the tower with minimal disturbance to nesting birds, and PVC tubes that facilitate delivery of whole thawed fish (capelin) 3 times per day at a subset of sites; see Gill and Hatch (2002) for details. All adults are banded, and nests are checked twice daily from initiation of nest building through fledging. Consequently, precise dates of egg laying, chick hatching and death/disappearance are known. The tower structure precludes predation by terrestrial and aerial predators, removing a major extrinsic source of variation in chick survival. On hatch day and every 5 days afterwards, all chicks are weighed, measured and marked on the head with a colored pen to indicate hatch order (facilitating identification in videos).

All nests used in this experiment contained two chicks and were tested in the morning that the first-hatched (hereafter “A chick”) was 5 days post-hatch. All second-hatched (hereafter “B chicks”) were 2 – 5 days post-hatch. All data were collected in June and July of 2021 (experiment 1) and 2022 (experiment 2), between 0545 and 0900 hrs, which was prior to any handling or research-related disturbance to the colony such as supplemental feeding or chick-measuring.

### 2.2 Supplemental feeding

Kittiwake chicks eat semi-digested food regurgitated by their parents. As part of a long-term experiment, a subset of tower nests were provided with supplemental food for the duration of the breeding season (hereafter “food supplemented” nests). Each food supplemented nest was equipped with a PVC pipe that allowed minimally-disruptive delivery of fish from inside the tower. All occupants of food supplemented nests were offered whole fish *ad libitum*, 3x per day, from the start of nest establishment through fledging, to augment the food they acquired at sea. Behavioral experiments were conducted when first-hatched chicks were 5 days old and too small to consume the supplemental fish directly, so any benefits of food supplementation were mediated by parents. However, by day 10 post-hatch they regularly consume the supplemental fish. “Control” chicks came from nests in which no supplemental food was provided, such that parents were entirely dependent on their own at-sea foraging to feed themselves and their chicks. This food supplementation paradigm successfully alleviates many nutritional constraints, more than doubling reproductive success in 2021 (control nests: ∼0.41 chicks per nest, versus ∼1.0 in food supplemented nests). For comparison, control nests in years with the highest reproductive success since the start of consistent monitoring (in 1996) produced ∼1.0 chicks per nest (Ferris and Zador, 2021).

### 2.3 EXPERIMENT 1. Endogenous corticosterone (Figure 1A)

In 2021, we collected video and chick blood samples from food supplemented (n = 16) and control (n = 17) nests. GoPro Hero Session 5 cameras were mounted on tripods inside the tower, and used to record each nest through the one-way mirrored glass for 62 ± 4.2 min. Blood sampling methods have been described previously (Benowitz-Fredericks et al., 2024) but in brief: At the end of the recording period, a timer was started, parents were alerted via a tap on the window, and the A chick was taken from the nest and blood-sampled from the alar vein using a sterile 26G needle and heparinized microhematocrit tubes. All samples were collected within 3 min of initial tap (range 46 - 120 sec, mean 93 ± 35 sec). We then placed the chick in a breathable cloth bag that we suspended 12 inches from a space heater to minimize thermoregulatory costs. Approximately 15 minutes (14 ± 1.8 min) from the time of original disturbance, the A chick was bled a second time and returned to the nest, followed by an additional 62 ± 6.5 min of video recording. High baseline corticosterone levels of undisturbed kittiwake chicks overlap with levels evoked by 15 min of restraint (Benowitz-Fredericks et al. 2024), suggesting that this protocol simulates ecologically-relevant levels of circulating corticosterone.

**Figure 1.**
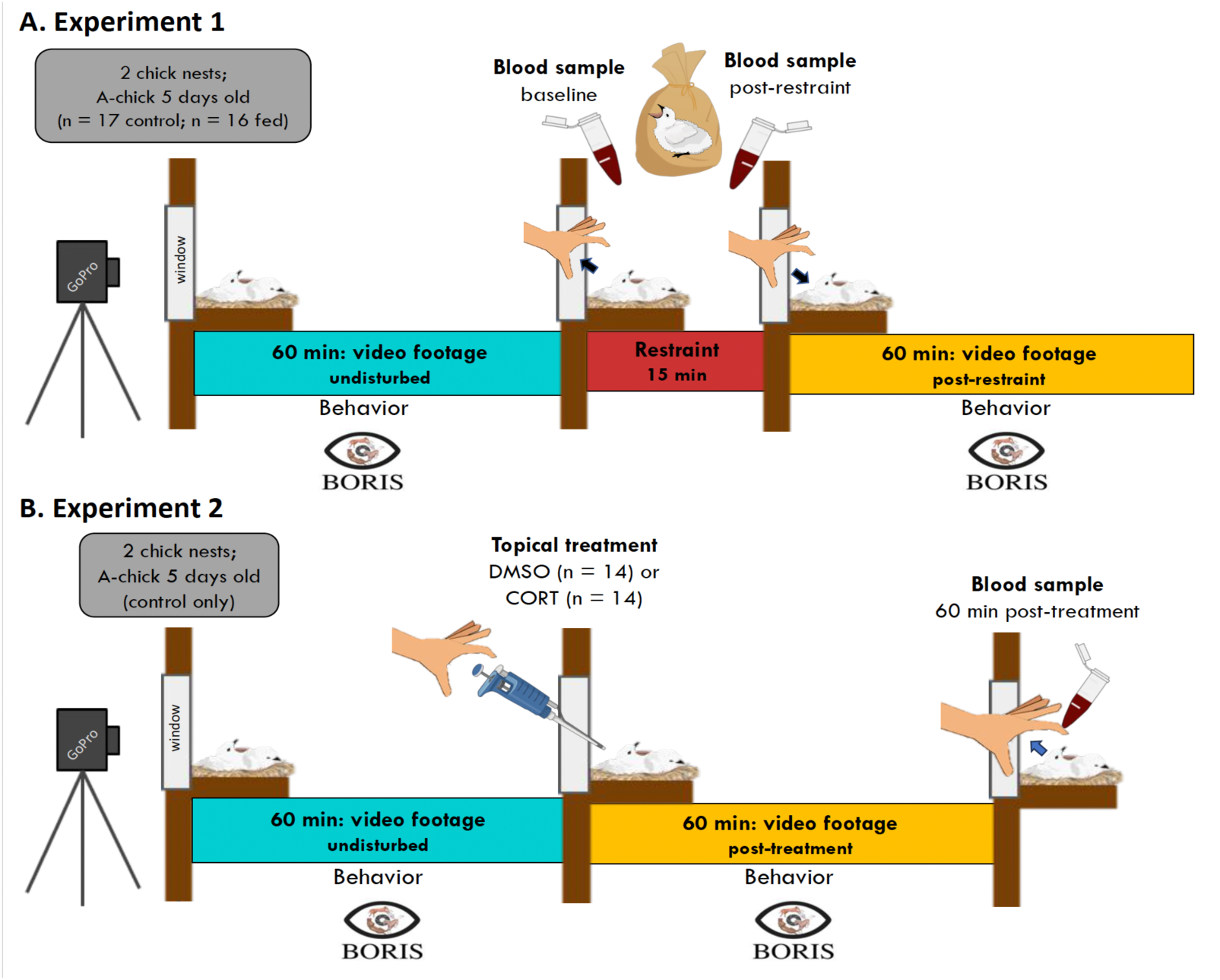
Experimental timelines. Timeline for behavioral recordings, treatments and blood sampling from 5-day old first-hatched chicks (“A-chicks”) in black-legged kittiwake nests containing two chicks. A) Experiment 1 (endogenous corticosterone). Undisturbed nests were video recorded for 60 min. Chicks were then bled in < 3min of nest disturbance, measured then restrained; the post-restraint bleed was taken 15 min after initial disturbance. Chicks were returned to the nest and recorded for another hour. B) Experiment 2 (exogenous corticosterone). Chicks were recorded for 60 min undisturbed, then briefly disturbed but not handled for topical treatment with DMSO or corticosterone in DMSO, recorded for another 60 min undisturbed, then captured for their first/only blood sampling at the end of the experiment. Some visual elements from BioRender.com

### 2.4 EXPERIMENT 2. Exogenous corticosterone (Figure 1B)

The second experiment was conducted in 2022 using only 5 day old A chicks from control (non-food supplemented) nests containing two chicks. The experimental design was similar to that in Experiment 1, however after 60.6 ± 1.3 min (mean ± SD) of video recording we administered minimally-invasive topical treatments of 12.4 ± 0.002µL of the veterinary solvent dimethyl sulfoxide (“DMSO”, n = 14), or of DMSO containing 5µg/µL of corticosterone (hereafter “CORT” treatment, n = 14). Because chick body mass varied, individual corticosterone doses varied, averaging 0.62 μg corticosterone per gram of body mass. Detailed methods for creating and administering the DMSO solutions, as well as the consequences for circulating corticosterone at 15 and 60 min following this topical treatment for kittiwake chicks, have been previously described (Benowitz-Fredericks et al., 2024). Briefly, we applied a drop of DMSO containing dissolved corticosterone (or DMSO alone, for the control group) to the crease at the nape of the neck of focal chicks without handling them or removing them from the nest. Following corticosterone administration, nests were video recorded for another hour (57.4 ± 5.6 min; mean ± SD) after which a single blood sample was taken from each A chick < 3 min after recapture from the nest (for genetic sexing and to measure corticosterone levels generated by topical treatments).

The effects of topical treatments on corticosterone have been published previously (Benowitz-Fredericks et al., 2024), but in short: samples taken from a different subset of chicks verified that within 15 min, this topical application elevated circulating corticosterone to levels similar to those evoked by the endogenous response to 15 min of handling and restraint. In focal chicks from this study, corticosterone was 6.4 ± 3.8 ng/mL in the DMSO treated chicks, and remained elevated at 27.7 ± 16.1 ng/mL in the corticosterone treated chicks 60 min after administration. Circulating corticosterone levels in the DMSO-treated control group 15 or 60 min after treatment did not differ from baseline levels in undisturbed chicks, confirming that the process of administering topical treatments did not itself evoke endogenous corticosterone elevation.

### 2.5 Hormone assays

After collection, blood was refrigerated until centrifugation for 5 minutes at ∼2000 rpm, after which plasma was separated from erythrocytes and both were frozen and transported at -20℃ for storage in the lab at -80℃. Samples from the same chick were run in the same assay; assays were conducted blind to chick sex or treatments. Corticosterone was extracted after adding tritiated corticosterone to calculate percent recovered (96% ± 3% (SD) and 92% ± 3% (SD) for experiments 1 and 2, respectively). 20 µL plasma samples were run in duplicate in two radioimmunoassays for experiment 1, and in a single assay for experiment 2, using an Esoterix (#B3-163) corticosterone antibody (lowest detectable limit: 7.8 pg/sample) following protocols described in Kitaysky et al. (2007). Final values were corrected for sample-specific percent recovery. Each assay included an internal corticosterone standard (inter assay CV for standard across three assays: 0.11%; intra assay CV for duplicates: 1%).

### 2.6 Genetic sexing

To genetically sex chicks, we used DNA that was extracted from frozen erythrocytes using Qiagen DNeasy kits, and PCR and gel electrophoresis to determine chromosomal sex of chicks using the primers and protocol described in Merkling et al. (2012).

### 2.7 Behavior analysis

Behavior of A chicks was coded using the software BORIS (Friard and Gamba, 2016) and a kittiwake chick ethogram (Appendix Table A1). Focal behaviors were aggression towards the B chick, begging, and feeding (calculated as bout rates - number of behavior bouts or events divided by total video time). To correct for variation among individuals in undisturbed behavior levels in both experiments, we quantified changes in behavior from the first to the second hour using residuals of the regression of behavior rates in the second hour on behavior rates in the first hour.

### 2.8 Data analysis

#### 2.8.1 Data preparation and models

To meet statistical assumptions for parametric analyses we log-transformed corticosterone concentrations if it improved normality. Thus corticosterone values were log_10_-transformed for experiment 1, but not for experiment 2. We also tested for equal variance between feeding treatment groups using the *var.test* function in R and accounted for unequal variance between feeding treatments in post-restraint corticosterone concentrations in all cases except when doing AICc model selection (see below), in which case we verified that the model output was the same for both approaches. We used generalized linear models with a Gaussian link except in the case of modeling survival, for which we used a binomial link, and untransformed corticosterone concentrations, for which we used an inverse gamma link. For outputs of top models, we report the effect size, a standardized coefficient with its 95% confidence interval (*effectsize*).

To determine what variable or combination of variables (models) best described the response we used Akaike’s Information Criteria corrected for small sample sizes (AICc) to rank models from a priori model sets that were built according to the hypotheses being tested. All models with a ΔAICc less than 2 were considered top performing and those with significant predictors are reported in the results. If the null model was included in the top performing model set, we considered the evidence for models with lower ΔAICc than the null as more viable than those ΔAICc values larger than the null, but considered the evidence for all models with ΔAICc <2 if the null model was not included in the top model set. We also report McFadden’s pseudo r-squared (values of McFadden’s pseudo-*R*^2^ between 0.2-0.4 are considered an excellent model fit, Hensher and Stopher, 2021) for generalized linear models (1-[residual deviance/null deviance]) and adjusted *R*^2^ as measures of goodness of fit. For all analyses, behavior changes were calculated using residuals of behavior bout rates from the first to the second hour. Finally, we checked for potential collinearity among closely related variables. In Experiment 1, baseline and post-restraint corticosterone were not correlated (F_1,31_ = 0.08, p = 0.77, adjusted *R*^2^ = -0.03), neither were pre-restraint aggression and begging rates (F_1,31_ = 2.84, p = 0.10, adjusted *R*^2^ = 0.05), nor pre- and post-restraint total feed rates (F_1,31_ = 0.17, p = 0.68, adjusted *R*^2^ = - 0.03). Pre- and post-restraint aggression (F_1,31_ = 6.28, p = 0.02, adjusted *R*^2^ = 0.14) and pre- and post-restraint begging rates (F_1,31_ = 4.09, p = 0.05, adjusted *R*^2^ = 0.087) were positively correlated, as were post-restraint aggression and begging rates (F_1,31_ = 9.69, p = 0.004, adjusted *R*^2^ = 0.21). In Experiment 2 pre- and post- total feed rates were not correlated (F_1,25_ = 0.16, p = 0.69, adjusted *R*^2^ = -0.03) while pre- and post- aggression rates (F_1,25_ = 19.39, p = 0.0002, adjusted *R*^2^ = 0.41) and pre- and post-begging rates (F_1,25_ = 8.29, p = 0.008, adjusted *R*^2^ = 0.22) were positively correlated.

#### 2.8.2 Experiment

Feeding treatment was a significant predictor of baseline corticosterone (t_30_ = 6.23, p < 0.001, higher in control nests) and our previous study showed that, as an interaction term with sex, it was an informative predictor of the HPA function (Benowitz-Fredericks et al. 2024). Therefore, we first ran the analyses with feeding treatment, sex, focal behaviors, and their interactions included (Appendix 2). We then re-ran the model selection analyses on control and supplementally fed chicks separately to disentangle the effects of feeding treatment on the results, given the constraints associated with the large number of predictor variables and our sample size, as well as our explicit interest in testing whether supplemental feeding weakened relationships between corticosterone levels and response variables (Hypothesis 3).

To test Hypothesis 1 (recent behaviors influence HPA-axis activity), we tested potential relationships between rates of behavior over the course of an hour and baseline and restraint-induced corticosterone at the end of the hour. For this analysis we ran separate model selections for baseline and post-restraint corticosterone, including models with begging, feeding and aggression bout rates from the hour of observation prior to blood sampling, as well as additive and interactive combinations of each behavior with sex.

To test Hypothesis 2, that elevated corticosterone affects behavior, we ran three model selection analyses with change in behavior (begging, aggression and feeding) as the response variable. We compared models that included baseline concentrations of corticosterone at the end of the first hour, concentrations of corticosterone elevation elicited by 15 min handling and restraint (endogenous stress response), and chick sex (as well as additive and interactive combinations) as predictors in models.

Finally, to evaluate potential fitness correlates of HPA activity (Hypothesis 4), we ran two model selection analyses, each of which included A chick baseline and restraint-induced corticosterone levels, and sex as potential predictor variables, as well as their additive and interactive combinations. We modeled A chick survival in response to their own baseline and restraint induced corticosterone levels, sex, feeding treatment and their interactions (we combined treatments for this analysis only as all chicks in the supplementary fed treatment survived, eliminating any variability in the response for the model to evaluate). We modeled the number of days the B chick survived in response to A chick baseline and restraint-induced corticosterone as well as sex and their interactions, analyzed separately by feeding treatment.

#### 2.8.3 Experiment 2

In the second experiment (exogenous corticosterone treatment), we compared changes in each behavior using residuals of behavior rates from the hour prior to treatment to the hour following treatment in both corticosterone-treated (hereafter “CORT”) and control chicks (hereafter “DMSO”) from non-supplemented nests only. We included both corticosterone treatment and corticosterone levels at the end of the experiment as predictor variables in our model set, because multicollinearity between the variables was moderate (variance inflation factor IF = 2.53), rendering it appropriate to include both as potential predictors of changes in behavior in the same statistical model.

## 2. RESULTS

### 3.1 Experiment 1

#### 3.1.1 Hypothesis 1:Behavior affects HPA-axis activity & Hypothesis 3: Relationships are modulated by food availability

In both supplemented and control nests we found no evidence for behavior, sex, or recent feeding experience as informative predictors of baseline corticosterone (Table 1, but see Table A3.1, for feeding treatment effect). For both feeding conditions the null model was the top performing model, with AICc values that were > 2 higher than the next best model (Table 1). We found evidence that post-restraint corticosterone in control, but not food supplemented nests, was positively correlated with the feeding rate in the previous hour (standard coefficient = 0.02 [0.01,0.03]; Table 1, Fig 2). Post-restraint corticosterone was also supported by an additive model with chick sex (feeding in previous hour + sex; sex effect: higher in males than females; standard coefficient = -0.02 [-0.04, 0.00]). In food supplemented nests, the null model was the top performing model (Table 1).

**Figure 2.**
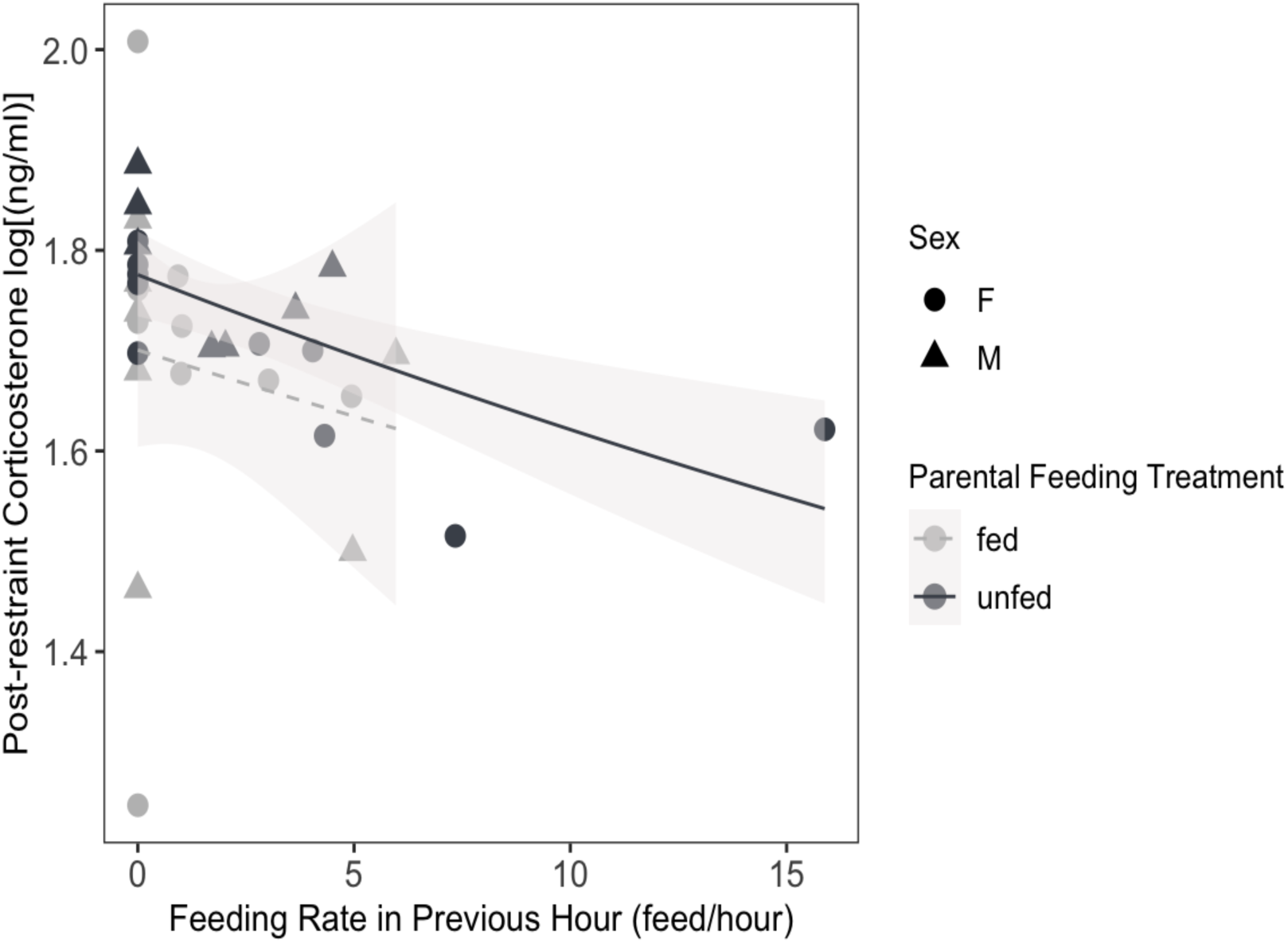
Feeding rates and subsequent restraint-induced corticosterone. In control nests where parents were not supplementally fed, corticosterone levels after 15 min of handling and restraint were higher in chicks who had received fewer feeds in the previous hour, and in male chicks. Dark lines = control feeding treatment, light lines = food supplemented treatment; solid lines = significant relationships and the dashed line = non-significant

**Table 1.** Model selection results for test of Hypothesis 1: behaviors drive corticosterone levels. Predictors in models included combinations of sex and behaviors in the previous hour. We ran separate model comparisons for chicks from control and food-supplemented nests. All models contributing to a cumulative weight of 95% are shown. Models within 2AIC units of the best performing model are indicated in bold, except for models with delta AICc values greater than the null model, which we we considered to be weakly supported. LL: log-likelihood, goodness of fit: McFadden’s pseudo r-squared (values between 0.2-0.4 are considered an excellent model fit).

| Response | Model | K | Delta AICc | AICc Wt | LL | Goodness of |
| --- | --- | --- | --- | --- | --- | --- |
| **Baseline corticosterone** |  |  |  |  |  |  |
| *Control* |  |  |  |  |  |  |
|  | **null** | **2** | **0** | **0.46** | **-3.18** | **0.000** |
|  | sex | 3 | 2.33 | 0.14 | -2.85 | 0.038 |
|  | begging previous hour | 3 | 2.97 | 0.1 | -3.17 | 0.001 |
|  | feeding previous hour | 3 | 2.98 | 0.1 | -3.17 | 0.000 |
|  | aggression previous hour | 3 | 2.99 | 0.1 | -3.18 | 0.000 |
|  | sex + begging previous hour | 4 | 5.8 | 0.03 | -2.84 | 0.039 |
| *Food supplemented* |  |  |  |  |  |  |
|  | **null** | **2** | **0** | **0.49** | **-1.05** | **0.000** |
|  | begging first hour | 3 | 2.95 | 0.11 | -0.99 | 0.007 |
|  | aggression previous hour | 3 | 3.03 | 0.11 | -1.03 | 0.003 |
|  | feeding previous hour | 3 | 3.04 | 0.11 | -1.03 | 0.002 |
|  | sex | 3 | 3.05 | 0.11 | -1.04 | 0.002 |
|  | sex + begging previous hour | 4 | 6.53 | 0.02 | -0.96 | 0.010 |
| **Post-restraint corticosterone** |  |  |  |  |  |  |
| *Control* |  |  |  |  |  |  |
|  | **feeding previous hour** | **3** | **0** | **0.46** | **21.98** | **0.451** |
|  | **sex + feeding previous hour** | **4** | **0.19** | **0.42** | **23.63** | **0.548** |
|  | sex * feeding previous hour | 5 | 4.12 | 0.06 | 23.72 | 0.553 |
| *Food supplemented* |  |  |  |  |  |  |
|  | **null** | **2** | **0** | **0.44** | **5.69** | **0.000** |
|  | aggression previous hour | 3 | 2.32 | 0.14 | 6.07 | 0.046 |
|  | begging previous hour | 3 | 2.62 | 0.12 | 5.92 | 0.028 |
|  | feeding previous hour | 3 | 2.66 | 0.12 | 5.9 | 0.025 |
|  | sex | 3 | 2.98 | 0.1 | 5.74 | 0.006 |
|  | sex + aggression previous hour | 4 | 5.92 | 0.02 | 6.09 | 0.048 |

#### 3.1.2 Hypothesis 2: Elevated corticosterone alters behavior & Hypothesis 3: Relationships are modulated by food availability *Aggression*

In control nests, we found strong evidence that sex and post-restraint corticosterone levels corresponded with changes in aggression, with both variables occurring in 3 of the 4 top-ranked models (Table 2; Fig 3A). The best-performing models were sex, the additive model of sex + post-restraint corticosterone, the model including a sex * post-restraint corticosterone interaction, and the model with post-restraint corticosterone only. Males had a larger increase in bouts of aggression post-restraint than females (standard coefficient = 0.16 [0.35, 1.96]). Higher levels of post-restraint corticosterone were positively correlated with increased aggression from the hour before to the hour after restraint (standard coefficient = 0.52 [0.08, 0.95]; Fig 3), this relationship was stronger in males (standard coefficient = 0.71 [0.10, 1.33]) than in females (standard coefficient = 0.26 [-0.41, 0.93]). In food supplemented nests, the null model was the top-ranked model (Table 2).

**Figure 3.**
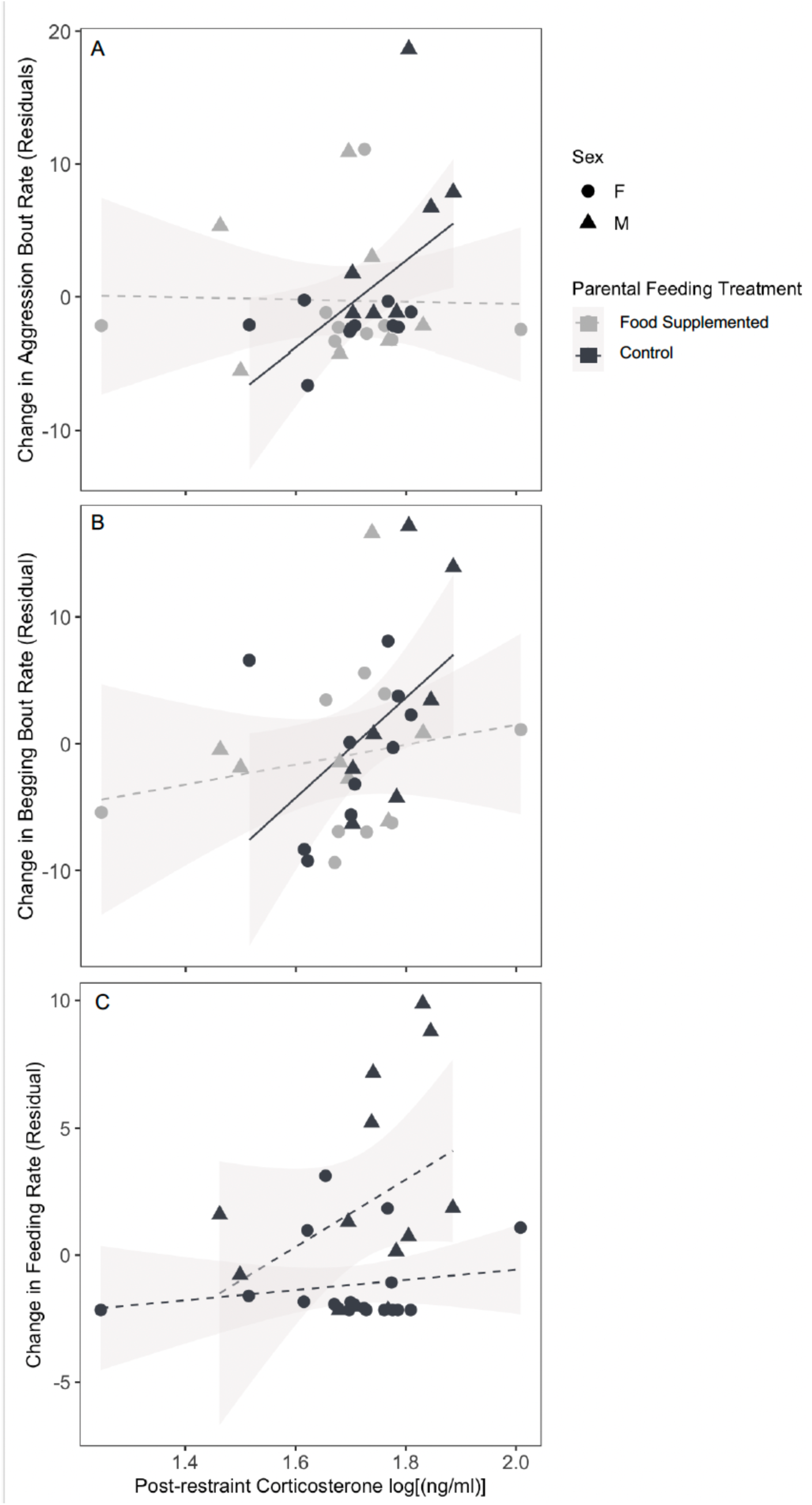
Restraint-induced corticosterone and subsequent changes in behavior. Changes in A. aggression, B. begging, and C. feeding. First-hatched kittiwake chicks with higher restraint-induced corticosterone levels subsequently showed larger increases in aggression and begging. However, these relationships were eliminated in experimentally food-supplemented nests. Behavior change was calculated as residuals from the regression of behavior bout rates (behavior bouts/hour) the hour before against behavior bout rates the hour after 15 min of handling and restraint (to control for interindividual variation in baseline behavior rates). Dark lines = control feeding treatment, light lines = food supplemented treatment; solid lines = significant relationships and the dashed line = non-significant.

**Table 2.** Model selection results for test of Hypothesis 2: elevated corticosterone redirects behavior. Response variables are the change in behavior from the pre- to post- endogenous corticosterone elevator. Predictors in models included combinations of sex, baseline corticosterone after the first hour of observation, and elevated corticosterone following 15 min of restraint, before the second hour of observation. We ran separate model comparisons for chicks from control and food-supplemented nests. All models contributing to a cumulative weight of 95% are included. Models within 2AIC units of the best performing model are indicated in bold, except for models with delta AICc values greater than the null model, which we considered to be weakly supported.LL: log-likelihood, goodness of fit: McFadden’s pseudo r-squared (values between 0.2-0.4 are considered an excellent model fit).

| Response | Model | K | Delta AICc | AICc Wt | LL | Goodness of Fit |
| --- | --- | --- | --- | --- | --- | --- |
| **Change in aggression** |  |  |  |  |  |  |
| *Control* |  |  |  |  |  |  |
|  | **sex** | **3** | **0** | **0.35** | **89.37** | **0.345** |
|  | **log post-restraint corticosterone + sex** | **4** | **1.31** | **0.18** | **90.46** | **0.423** |
|  | **log post-restraint corticosterone \* sex** | **5** | **1.74** | **0.14** | **92.31** | **0.536** |
|  | **log post-restraint corticosterone** | **3** | **1.93** | **0.13** | **88.41** | **0.266** |
|  | log baseline corticosterone + sex | 4 | 3.16 | 0.07 | 89.54 | 0.357 |
|  | null | 2 | 4.19 | 0.04 | 85.78 | 0.000 |
|  | log baseline corticosterone + log post-restraint corticosterone | 4 | 5.11 | 0.03 | 88.56 | 0.279 |
| *Food supplemented* |  |  |  |  |  |  |
|  | **null** | **2** | **0** | **0.32** | **82.69** | **0.000** |
|  | log baseline corticosterone | 3 | 0.46 | 0.26 | 84 | 0.151 |
|  | log baseline corticosterone + log post-restraint corticosterone | 4 | 1.18 | 0.18 | 85.46 | 0.293 |
|  | sex | 3 | 2.7 | 0.08 | 82.88 | 0.023 |
|  | log post-restraint corticosterone | 3 | 3.07 | 0.07 | 82.7 | 0.001 |
| **Change in begging** |  |  |  |  |  |  |
| *Control* |  |  |  |  |  |  |
|  | **log post-restraint corticosterone** | **3** | **0** | **0.43** | **83.96** | **0.238** |
|  | **null** | **2** | **1.64** | **0.19** | **81.64** | **0.000** |
|  | sex | 3 | 3.41 | 0.08 | 82.25 | 0.069 |
|  | log baseline corticosterone + log post-restraint corticosterone | 4 | 3.43 | 0.08 | 83.98 | 0.241 |
|  | log post-restraint corticosterone + sex | 4 | 3.44 | 0.08 | 83.98 | 0.241 |
|  | log post-restraint corticosterone * sex | 5 | 4.12 | 0.06 | 85.7 | 0.380 |
| *Food supplemented* |  |  |  |  |  |  |
|  | **null** | **2** | **0** | **0.48** | **79.02** | **0.000** |
|  | sex | 3 | 2.14 | 0.16 | 79.48 | 0.057 |
|  | log post-restraint corticosterone | 3 | 2.38 | 0.14 | 79.36 | 0.042 |
|  | log baseline corticosterone | 3 | 2.99 | 0.11 | 79.06 | 0.006 |
|  | log post-restraint corticosterone + sex | 4 | 4.9 | 0.04 | 79.93 | 0.107 |
| **Change in feeding** |  |  |  |  |  |  |
| *Control* |  |  |  |  |  |  |
|  | **sex** | **3** | **0** | **0.44** | **97.86** | **0.269** |
|  | null | 2 | 2.35 | 0.14 | 95.19 | 0.000 |
|  | log post-restraint corticosterone | 3 | 2.7 | 0.11 | 96.51 | 0.144 |
|  | log post-restraint corticosterone + sex | 4 | 2.86 | 0.11 | 98.17 | 0.270 |
|  | log baseline corticosterone + sex | 4 | 3.48 | 0.08 | 97.86 | 0.296 |
|  | log post-restraint corticosterone * sex | 5 | 4.4 | 0.05 | 99.46 | 0.395 |
| *Food supplemented* |  |  |  |  |  |  |
|  | **sex** | **3** | **0** | **0.31** | **90.76** | **0.190** |
|  | **null** | **2** | **0.23** | **0.28** | **89.1** | **0.000** |
|  | log post-restraint corticosterone + sex | 4 | 1.96 | 0.12 | 91.59 | 0.267 |
|  | log post-restraint corticosterone | 3 | 2.28 | 0.1 | 89.62 | 0.062 |
|  | log baseline corticosterone | 3 | 3.2 | 0.06 | 89.16 | 0.007 |
|  | log baseline corticosterone + sex | 4 | 3.56 | 0.05 | 90.79 | 0.190 |

##### Begging

In control nests, changes in begging bout rate were best described by post-restraint corticosterone, with the null as the next best model. Higher post-restraint corticosterone was positively correlated with an increase in begging bout rate (standard coefficient = 0.49 [0.05, 0.93]; Table 2, Fig 3B). The null model was the highest ranked model for chicks from food supplemented nests (Table 2).

##### Feeding

Changes in feeding rate were best described by chick sex, with the null model as the second-best model, in both control (standard coefficient = 1.02 [0.17, 1.88]) and food supplemented nests (standard coefficient = 0.84 [-0.08, 1.77], Table 2). Males had larger increases in their feeding rates than females (Fig 3C). Post-restraint corticosterone only appeared in top models when feeding treatments were combined (Appendix Table A2.5).

#### 3.1.3 Hypothesis 4: HPA-axis activity has fitness-related implications

Five out of 18 A chicks from control nests (without food supplementation) didn’t survive to fledging, while all 16 food supplemented A chicks survived; the ratio of surviving A chicks tended to be lower in the non-supplemented treatment (Yates corrected Chi-square=3.23, 1 d.f., p= 0.072). Because so few A chicks died, sample sizes were not sufficient for an analysis of A chick survival duration. However, survival of A chicks was best explained by baseline corticosterone and feeding treatment. The null model was also included in the top model set (Table 3). A chicks with higher baseline corticosterone were more likely to die than those with lower levels of baseline corticosterone; for each 1 ng/ml increase in baseline corticosterone the probability of chick survival decreased by 9% (odds ratio: 0.92, 95% confidence interval [0.81, 1.00]). A chicks that survived had, on average, lower baseline corticosterone (8.04 ng/mL ±8.01 (SD) vs 19.83 ng/mL ±16.2 (SD) for live vs dead, correspondingly, beta (ß)=-0.415 [-0.743, -0.087]).

**Table 3.**
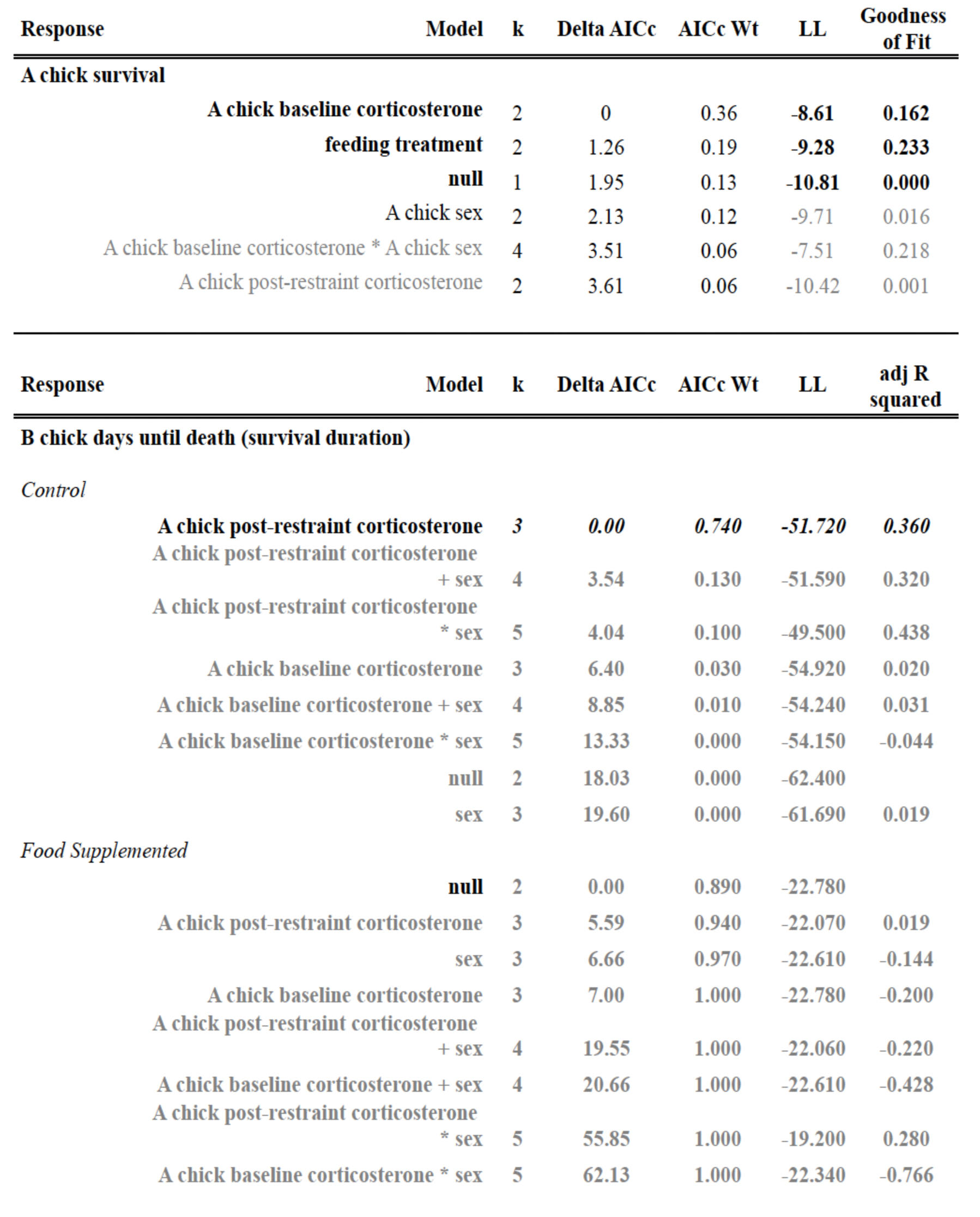
Model selection results for tests of Hypothesis 4: survival of A and B chicks, considering sex and endogenous corticosterone levels of the A chick. Models with delta AICc < 2 are in bold. All models contributing to a cumulative weight of 95% are included. LL: log-likelihood, goodness of fit: McFadden’s pseudo r-squared (values between 0.2-0.4 are considered an excellent model fit).

In control nests, A chick post-restraint corticosterone negatively correlated with the length of time B chicks survived post experiment (Fig. 4; standard coefficient = -0.64 [-1.10, -0.18]); there was no correlation in supplementally fed nests (standard coefficient = 0.43 [-0.61, 1.47]). 15 out of 18 control B chicks and 7 out of 16 food supplemented B chicks did not survive to fledging; the ratio of surviving B chicks was lower in control treatment (Yates corrected Chi-square=4.21, 1 d.f., p=0.04).

**Figure 4.**
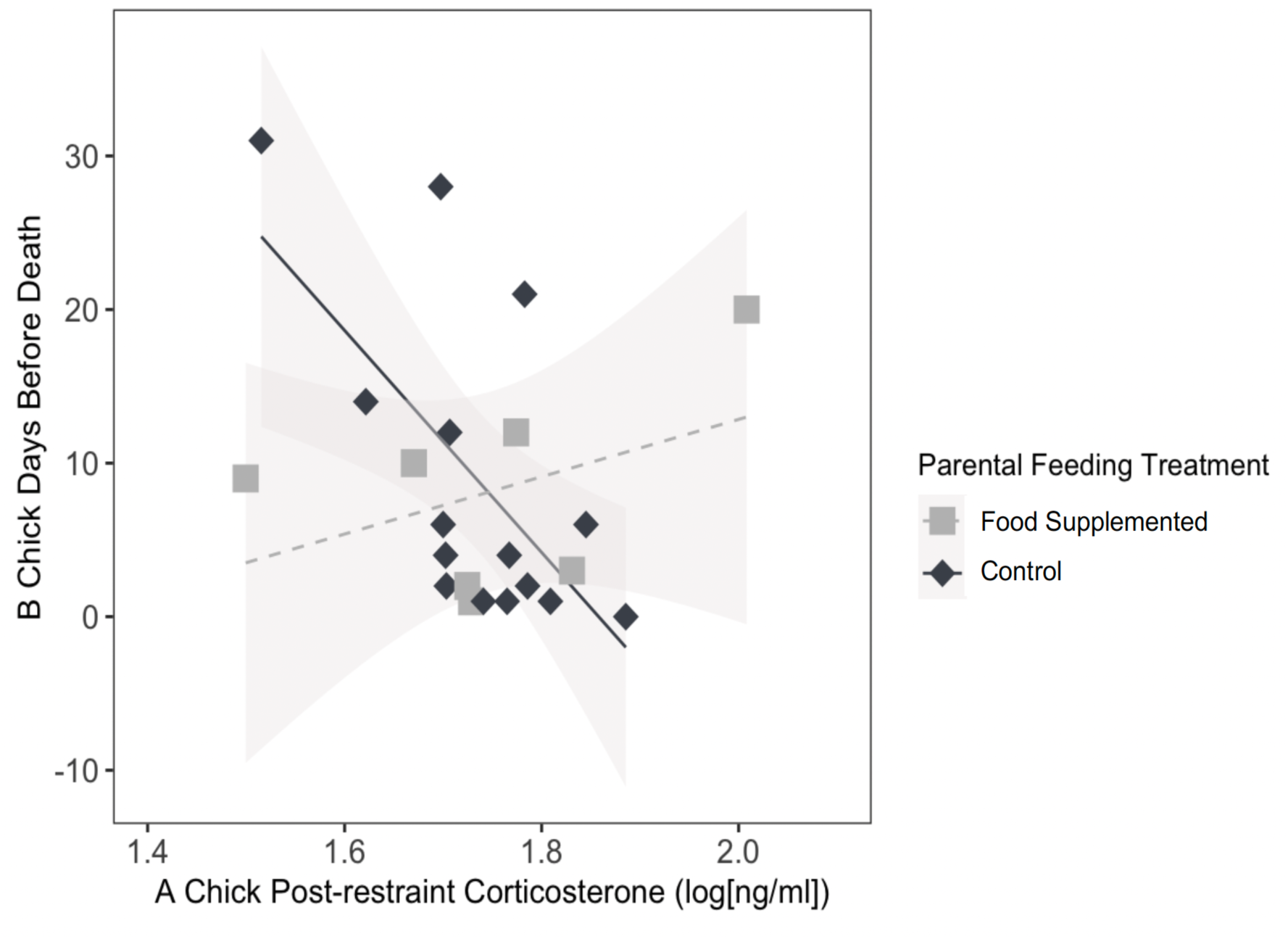
A chick corticosterone and B chick survival duration. B chicks in the control (no supplemental feeding) treatment died sooner when their 5-day old A chick siblings had higher levels of endogenous corticosterone after 15 min of restraint. Line darkness indicates treatment, solid lines = significant relationships and the dashed line = non-significant.

### 3.2 Experiment 2: Testing hypothesis 2 with exogenous corticosterone

When we looked at the effect of experimentally elevated corticosterone on behavior in chicks from non-supplemented nests, we found no evidence that exogenously applied corticosterone induced changes in begging bout rates or feeding rates - none of our models for begging or feeding rate were ranked above the null model (Table 4).

**Table 4.** Model selection results for experiment 2 (changes in behavior from 1 hour before to 1 hour after exogenous corticosterone elevation using topical treatments). Corticosterone treatments were topical DMSO (control), or corticosterone in DMSO admi.nistred to 5 day old A chicks. Chick corticosterone was measured once at the end of the experiment, 1 hour after topical treatments. All chicks were from non-food supplemented nests. All models contributing to a cumulative weight of 95% are included. Models within 2 AIC units of the best performing model are indicated in bold. LL: log-likelihood, goodness of fit: McFadden’s pseudo r-squared (values between 0.2-0.4 a1e considered an excellent model fit).

| Response | Model | K | Delta AICc | AICc Wt | LL | Goodness of Fit |
| --- | --- | --- | --- | --- | --- | --- |
| **Change in aggression** |  |  |  |  |  |  |
|  | **corticosterone treatment** | **3** | **0** | **0.24** | **130.62** | **0.138** |
|  | **post experiment corticosterone \* sex** | **5** | **1.3** | **0.12** | **130.27** | **0.271** |
|  | **post experiment corticosterone** | **3** | **1.33** | **0.12** | **131.27** | **0.095** |
|  | **sex + corticosterone treatment** | **4** | **1.47** | **0.11** | **128.61** | **0.179** |
|  | **null** | **2** | **1.47** | **0.11** | **130.97** | **0.000** |
|  | post experiment corticosterone + sex | 4 | 2.26 | 0.08 | 130.72 | 0.154 |
|  | post experiment corticosterone * corticosterone treatment | 4 | 2.59 | 0.07 | 129.26 | 0.145 |
|  | sex | 3 | 2.72 | 0.06 | 131.53 | 0.047 |
|  | sex + corticosterone treatment + post experiment corticosterone | 5 | 4.11 | 0.03 | 131.44 | 0.191 |
| **Change in begging** |  |  |  |  |  |  |
|  | **null** | **2** | **0** | **0.34** | **131.52** | **0.000** |
|  | **post experiment corticosterone** | **3** | **0.74** | **0.23** | **132.43** | **0.065** |
|  | corticosterone treatment | 3 | 2.29 | 0.11 | 131.65 | 0.009 |
|  | sex | 3 | 2.54 | 0.09 | 131.52 | 0.000 |
|  | post experiment corticosterone + corticosterone treatment | 4 | 3.21 | 0.07 | 132.58 | 0.075 |
|  | post experiment corticosterone + sex | 4 | 3.5 | 0.06 | 132.43 | 0.065 |
|  | post experiment corticosterone * corticosterone treatment | 5 | 4.89 | 0.03 | 133.26 | 0.121 |
| **Change in feeding** |  |  |  |  |  |  |
|  | **null** | **2** | **0** | **0.39** | **148.66** | **0.000** |
|  | post experiment corticosterone | 3 | 2.11 | 0.13 | 148.88 | 0.016 |
|  | corticosterone treatment | 3 | 2.26 | 0.12 | 148.8 | 0.010 |
|  | sex | 3 | 2.42 | 0.12 | 148.72 | 0.005 |
|  | post experiment corticosterone + corticosterone treatment | 4 | 3.06 | 0.08 | 149.79 | 0.080 |
|  | post experiment corticosterone + sex | 4 | 4.71 | 0.04 | 148.97 | 0.022 |
|  | sex + corticosterone treatment | 4 | 4.89 | 0.03 | 148.87 | 0.016 |
|  | post experiment corticosterone * corticosterone treatment | 5 | 5.17 | 0.03 | 150.25 | 0.111 |

However, we found strong evidence that exogenous corticosterone treatment induced an increase in aggression: changes in aggression bout rates were lower in chicks treated with DMSO alone compared to those treated with exogenous corticosterone (standard coefficient = -0.73 [-1.44, -0.02]; Table 4). Sex and corticosterone levels 60 min post-treatment also contributed to explaining changes in aggression, with both variables occurring in 3 of the 4 top-ranked models (Table 4). One of these models was the interactive model of sex * post-treatment corticosterone (Table 4, Fig. 5), revealing that corticosterone levels at the end of the hour were positively correlated with an increase of aggression bout rates in males (standard coefficient = 0.67, [0.24,1.11]) but not in females (standard coefficient = 0.12, [-0.44, 0.68]).

**Figure 5.**
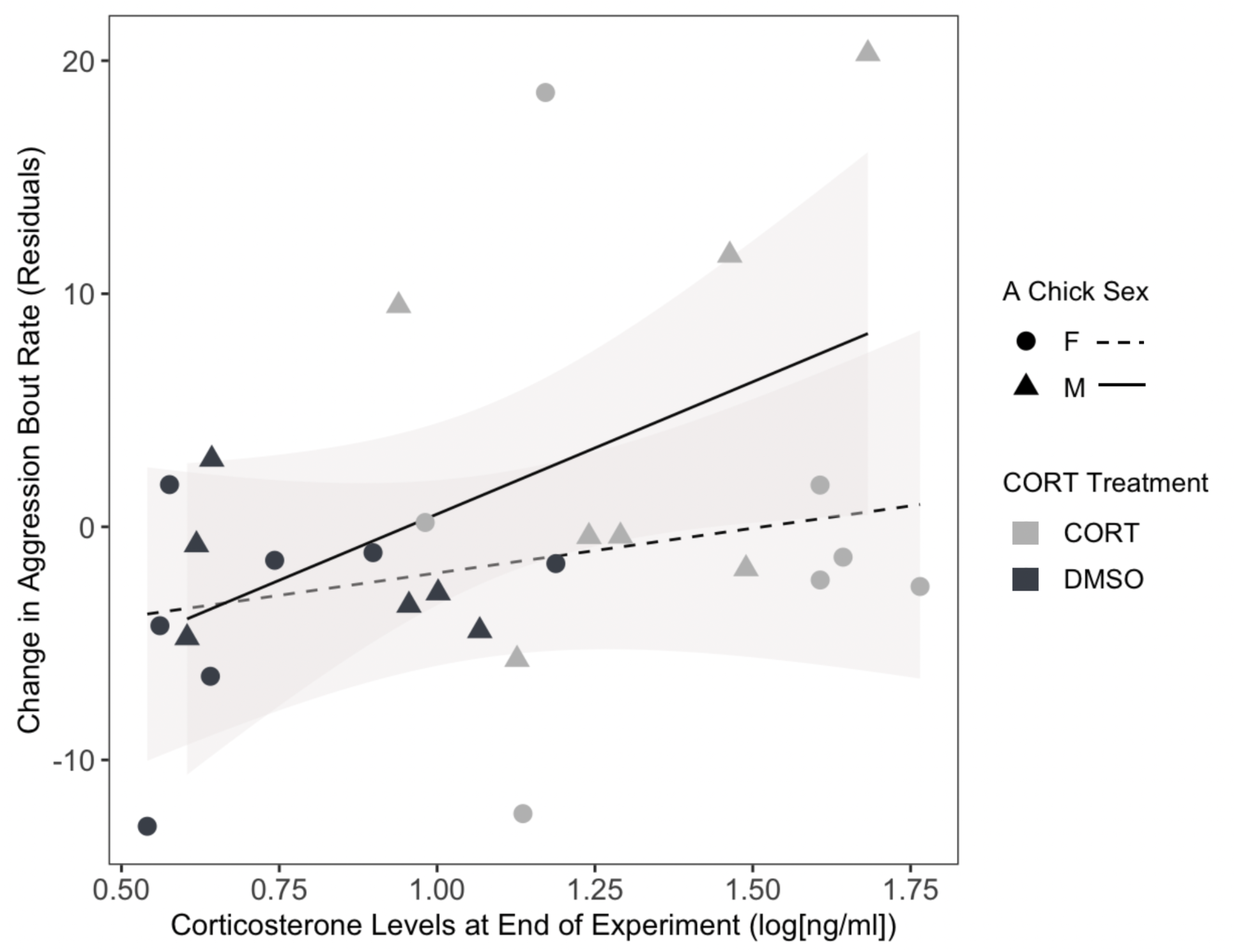
Exogenously-elevated corticosterone levels and changes in aggression. Male (but not female) chicks from nests without food supplementation that had higher corticosterone levels 60 min after treatment with topical corticosterone had larger increases in aggression, calculated as residuals from the regression of aggression bout rates (bouts/hour) the hour after treatment on aggression bout rates the hour before treatment (to control for interindividual variation in baseline behavior rates). Solid line = male; dashed line = female.

## 3. DISCUSSION

### 4.1 Overview

We found some form of support for each of the four hypotheses we tested. While we did not find strong evidence that behavior drives baseline HPA-axis activity in young kittiwakes (Hypothesis 1), we did find that post-restraint corticosterone levels were higher in chicks who had received fewer feeds in the previous hour (but not in chicks who had engaged in more begging or aggression). In contrast, Hypothesis 2 was well-supported: post-restraint corticosterone levels were positively associated with subsequent changes in behavior (increases in begging in both sexes, and aggression in males). The second experiment, in which we administered topical corticosterone, further supported Hypothesis 2 and suggested that the corticosterone:aggression relationship is the one most likely to be causal. The increase in begging was not retained when corticosterone was elevated in isolation, and may have been caused or modulated by other components of the stress response (such as catecholamines; see section 4.5 below). Hypothesis 3 was also well-supported: most of the hormone:behavior relationships were eliminated in nests where food limitation was alleviated via supplemental feeding. Finally, the magnitude circulating corticosterone after 15 min of restraint was negatively associated with the duration of survival of younger siblings, providing support for fitness relevance of HPA axis activity (hypothesis 4). Together, these results support the hypothesis that elevation of corticosterone can rapidly influence behavior in context- and sex- specific ways. In the rest of the discussion, we focus on the results that we find most interesting and important.

### 4.2 Sex differences

Though chromosomal sex imperfectly captures sources of interindividual variation (Smiley et al. 2024), in this study it was effective at explaining some of the variation in hormone:behavior relationships. There are sex differences in kittiwake chick growth rates and survival, often attributed to preferential parental allocation (Merkling et al., 2012; Young et al., 2016), however sex differences in the relationships between chick behavior and HPA axis activity have not been studied. In fact, we were only able to identify one study that considered (but did not find) sex-specific relationships between corticosterone and chick behavior in any bird (the white-crowned sparrows; Wada and Breuner, 2008). Our study suggests that behavioral strategies for coping with stressors may be sex-specific in 5 day old kittiwake chicks, despite a lack of consistent dimorphism in morphology or HPA axis activity at this age (Benowitz-Fredericks et al., 2024). Post-restraint and exogenously elevated corticosterone levels were positively correlated with changes in aggression in males but not females, though body size does not diverge until males become larger and heavier around 15 days post hatch; Merkling et al., 2012). We previously showed that sex did not affect corticosterone levels at baseline or in response to 15 min of handling and restraint in 5 day old kittiwake chicks in the years of this study (Benowitz-Fredericks et al., 2024). However, there may be underlying sex differences in molecular substrates like glucocorticoid receptor expression (Jimeno et al., 2023), which can vary without concurrent differences in HPA axis activity (Cornelius et al., 2018), such that relationships between corticosterone and behavior may still vary by sex even if circulating levels of corticosterone do not. Given that sex-specific strategies for maximizing fitness can include differences in both developmental rates (Richner, 1991; Weimersckirch et al., 2000) and somatic state at the end of development (Metcalfe and Monaghan, 2001), our results suggest that female chicks experiencing elevated corticosterone may be able to maximize their fitness without escalating competitive behaviors.

### 4.3 Magnitude of restraint-induced corticosterone is positively related to changes in behavior

Our study supports the overarching hypothesis that elevated corticosterone can rapidly increase food acquisition behaviors in chicks.

#### 4.3.1 Aggression

In both experiments, we found a positive relationship between aggression and elevated corticosterone (though not baseline corticosterone) that was stronger in male chicks compared to females. To our knowledge, a single study in birds has looked at relationships between corticosterone and aggression on acute time scales, but asked whether aggressive behavior influenced corticosterone levels, rather than whether corticosterone influenced aggressive behavior. Consistent with our results that recent aggression did not affect baseline corticosterone, a study of siblicidal Nazca booby chicks did not find any changes in corticosterone immediately following major aggression bouts (Ferree et al. 2004). The effects of glucocorticoids on aggression have been better-studied in mammals than in birds, and such studies generally conclude that the relationship depends on the social context and the nature of glucocorticoid elevation (acute, chronic, or “toxic”; Haller, 2020, Yohe et al., 2012). Our study provides strong evidence that corticosterone may have a causal role in rapidly increasing context-appropriate aggression in very young kittiwake chicks.

#### 4.3.2 Begging

There is variation among species and studies in the amount of energy expended during begging, but there is a general agreement that begging does not contribute substantially to most nestlings’ daily energy budget (Morena-Rueda, 2007). The relatively low metabolic cost of begging may explain the lack of relationship between begging rates and subsequent baseline corticosterone levels in our study. A handful of studies have tested whether elevated corticosterone is associated with variation in begging behavior in free-living passerine chicks on the scale of minutes, usually administering a series of corticosterone treatments over multiple days and finding relatively small changes (Bowers et al. 2019, Elderbrock et al., 2017, Wada and Breuner, 2008). We found stronger positive relationships between increases in begging and the magnitude of endogenous corticosterone following restraint than between begging and circulating corticosterone following exogenous corticosterone administration. This suggests either that the begging:corticosterone relationship may be due to other components of the endogenous stress response that correlate with corticosterone, or that there is a behavior-specific, dose-dependent effect of corticosterone on begging behavior (see section 4.5).

#### 4.3.3 Feeding

Feeding rates by parents had no effect on baseline corticosterone. While frequent or recent feeding might be predicted to reduce corticosterone levels as net energy balance improves, postprandial elevations in glucocorticoids (presumably associated with changes in intermediary metabolism) may confound this effect. Post-prandial glucocorticoid elevations have been well-documented on the scale of 30-60 min in humans (Slag et al., 1981, Anderson et al., 2019) and in chicks of at least one species of bird (the Florida scrub jay; Elderbrock et al., 2018). In contrast to baseline levels, post-restraint corticosterone levels were negatively correlated with feeds in the previous hour, suggesting increased reactivity of the HPA axes of chicks who had received less food recently. It is unclear whether this was due to the events of the past hour, or due to a correlation between feeding rates in our hour of observation and long-term feeding rates, though changes in HPA axis responsiveness are thought to reflect an individual’s recent history of HPA axis activation (Dallman, 1993). Post-restraint corticosterone levels were not related to subsequent feeding when control and food supplemented nests were analyzed separately, potentially because feeding is moderated by parental responsiveness to chick behavior, rather than being controlled by chicks. However, when feeding treatments were combined for model analysis, post-restraint corticosterone emerged in two of the three top models explaining subsequent feeding rates (Appendix 2, Table A.2.5), suggesting that while the relationship is weaker than for aggression or begging, it is present.

### 4.4 Rapid behavioral effects of corticosterone

An unusual aspect of our study is that it focuses on glucocorticoid:behavior relationship on the scale of an hour, rather than days or weeks. We found evidence that corticosterone may be able to facilitate behavioral changes on this short time scale. In seabirds, prolonged elevation of exogenous corticosterone in chicks tends to increase begging rates on the scale of days (Kidawa et al., 2017; Kitaysky et al., 2001; Kitaysky et al., 2003; Loiseau et al., 2007; Vallarino et al. 2006). However, the HPA axis is also highly responsive in young chicks of many species, capable of substantially elevating glucocorticoids on the scale of minutes (Bebus et al., 2020). While rapid effects of glucocorticoids on behavior are not consistent with classical genomic mechanisms of action for glucocorticoid receptors, putative membrane glucocorticoid receptors in birds may allow rapid non-genomic effects of corticosterone (Breuner and Orchinik, 2009; Gao and Deviche, 2018). There is also evidence in mammals that “classical” nuclear steroid receptors can be found outside the nucleus, where they evoke rapid, non-genomic responses (Ordóñez-Morán and Muñoz 2009; Wilkenfeld et al. 2018). The accumulating body of evidence that corticosterone can evoke rapid effects in birds (Breuner et al., 1998, this study) and the conserved steroid hormone related pathways across vertebrates suggest that birds are likely to share such mechanisms of action for glucocorticoids. This evidence, combined with growing interest in rapid increases in corticosterone and their consequences (Taff et al., 2022), suggests that the phenomenon warrants continued investigation across species and ages.

### 4.5 Other components of the stress response?

The endogenous response to stressors include a suite of physiological changes (Gaidica & Dantzer, 2020; Gormally & Romero, 2020), of which elevated glucocorticoids is just one. The inability to separate the effects of corticosterone from the effects of other components of the stress response like catecholamines makes it challenging to interpret relationships between experimentally-induced increases in endogenous corticosterone and other physiological and behavioral variables of interest. Our second experiment, using minimally disruptive delivery of exogenous corticosterone, isolated potential effects of corticosterone, allowing us to attribute causality to the relationship between corticosterone levels and aggression that we found in experiment 1. Our results do not necessarily exclude the possibility that begging and feeding are also driven by elevated corticosterone for three reasons. First, experiment 1 and 2 were conducted in two different years, and interannual variability in chick or parental physiological state could generate variation in responses to elevated corticosterone. Second, behavioral responses may change along a continuum of corticosterone levels, triggered by dose dependent activation of different receptors (Mifsud & Ruel, 2018; Proszkowiec-Weglarz & Porter, 2010), with aggression being facilitated by lower levels of corticosterone than begging and feeding. While topical treatment significantly elevated corticosterone, average levels 15 min after treatment were lower in the exogenous CORT treatment than those generated by the endogenous response to 15 min of restraint (Benowitz-Fredericks et al. 2024). Finally, our exogenous treatment kept corticosterone elevated above baseline for at least 60 min but we do not know how long endogenous corticosterone remained elevated following the 15 min restraint protocol. It is possible that sustained corticosterone elevations affect the three focal behaviors differently than ephemeral corticosterone elevations.

### 4.6 Challenges reveal relationships

Our study provided two lines of evidence that, as seen in adult kittiwakes (Benowitz-Fredericks et al., 2022), relationships between behavior and corticosterone in chicks emerge in response to challenges. First, baseline corticosterone levels were not strongly related to behaviors before or after the sample was taken - hormone:behavior relationships emerged after chicks were exposed to a stressor (handling and bag restraint) or experimental manipulation (exogenous corticosterone) that elevated corticosterone. Killen et al. (2013) described the potential role of environmental challenges in shaping the emergence of relationships between physiology and behavior, proposing that they can either attenuate/ mask them, or amplify them, as we found in this study. Second, supplemental feeding eliminated most of the relationships between endogenous corticosterone and behavior. Interestingly, chicks from food supplemented nests reached the same average levels of corticosterone as chicks from non-supplemented nests after 15 min of bag restraint (secreting more corticosterone to do so, since they started at lower baseline levels; Benowitz-Fredericks et al. 2024). Thus, the elimination of hormone:behavior relationships in food-supplemented nests was not due to a reduction in HPA sensitivity or capacity. Instead, our results are consistent with the basic tenet of hormone:behavior relationships that hormones do not cause behaviors, but increase the likelihood of their expression in response to appropriate stimuli. Chicks from food supplemented nests were less likely to be in a hungry state, therefore it was less likely that an elevation in corticosterone would evoke increases in behaviors associated with hunger (begging and aggression).

### 4.7 HPA axis capacity and fitness

The endocrine physiology of A chicks weakly negatively predicted their own survival, and strongly negatively predicted survival duration of their younger siblings. In experiment 1, nearly all of the A chicks survived, though those that died had higher baseline corticosterone than those that survived. Most of the B chicks died, which is normal for non-supplemented nests at this colony, where average fledging success is <1 chick per nest (Ferris and Zador, 2021). However, in nests where B chicks died, they did so more quickly when they had A chick siblings who mounted larger corticosterone responses to restraint. More rapid death of B chicks with A siblings who had greater HPA responsiveness could be directly related to the phenotype of the A chick (since chicks with more responsive HPA axes were more aggressive), though the proximate cause of death can rarely be determined in the field. Alternatively, it could be due to the fact that in nests where parents were low quality or otherwise struggling to sufficiently provision chicks, both A chick HPA axis reactivity and B chick survival reflect lower provisioning rates by parents.

The fitness impacts of B chick deaths for A chicks are nuanced, since siblicide reduces indirect fitness but, presumably, increases direct fitness. Elimination (and more rapid elimination) of competition for parental provisioning should increase a chick’s energy budget by increasing food intake and potentially reducing expenditure for competitive behaviors like aggression. Brood-size manipulations in kittiwakes support this assumption, finding increased growth and survival of A chicks in reduced broods compared to control or enlarged broods (Jacobsen et al., 1995; Young et al., 2017). Our study suggests that siblicidal “decisions” in this species with facultative siblicide species may be facilitated by corticosterone.

### 4.8 Conclusion

The ability of very young, nest-bound chicks to release large quantities of corticosterone despite the potential costs of exposure to corticosterone during development suggests that such an early onset of HPA-axis responsiveness to nutritional state and external stimuli should increase their fitness despite their somewhat limited ability to change their environment. Our finding that when food is limited, the magnitude of corticosterone released in response to a challenge is correlated with subsequent increases in a repertoire of behaviors that promotes resource acquisition and brood reductions is consistent with this hypothesis. In addition, the emergence of sex-specific relationships between corticosterone release and behavior at an age where other sex differences are undetectable suggests the existence of sex-specific differences in proximate molecular mechanisms (such as the location, density, and/or downstream paths for glucocorticoid receptors) and ultimate differences in male and female behavioral strategies during development. To better understand the selection pressures shaping patterns of HPA axis ontogeny across species, we encourage future research about relationships between glucocorticoids and behavior in free-living juvenile animals to consider: a) effects on short time-scales, b) the potential for sex-specific strategies, and c) the role of environmental challenges in masking or revealing potential relationships among them. Finally, glucocorticoids are fundamentally metabolic hormones, so understanding the rapid metabolic correlates and consequences of corticosterone elevation in young chicks will further elucidate the function of HPA axis activity in this context.

## Acknowledgements

We are grateful to the following - Eadaoin Kelly for indispensable assistance in the field and Fred Tremblay, Kyle Elliott, and Scott and Martha Hatch for field management and logistical support. For months of feeding and monitoring kittiwake nests: Catherine Lee-Zuck, Jess Morales-Valenzuela, Kelly Bacile, Julia Dunoyer, Aidan Colligan and Eliza Fernandes-McDade (2021) and Gabby Dennis, Hailey Gee, Lauren Jackson, David Jahdon, Alex Turmaine, Léo Marcouillier, Chinatsu Nakajima, and Stella Solasz (2022). Ali Jackson and Alex Le: genetic sexing and Taiba Khan: help with ethogram development. Two anonymous reviewers provided helpful feedback on the manuscript. This project was funded by a Gates Millennial Fellowship to SNP, Bucknell Dean’s Fellowship to ZMBF, Gulf Watch Alaska and Alaska Ocean Observing System (SW/Institute for Seabird Research and Conservation), and the Institute of Arctic Biology, University of Alaska Fairbanks (ASK and APW). All research was conducted under permits from the U.S. Fish and Wildlife Service (#MB33779D-1), Alaska Department of Fish and Game (#21-089) and with approval from McGill University’s Animal Use Committee.

## Appendix A1. Ethogram of chick behaviors

| **Behavior** | **Point/State** | **Behavior Definition** | **Bout Definition** |
| --- | --- | --- | --- |
| Begging | State | 1. The chick's beak is oriented upwards or in the direction of the parent's beak paired with a thrusting/lunging motion of head toward parent's beak. Sometimes the chick also opens and closes beak<br>2. Thrusting varies depending on the distance of the chick's beak from the parent's beak.<br>3. Can be accompanied by peeping.<br>4. A typical behavior before, during, and in between begging bouts is the chick's head constantly following the pathway of the parent's head/beak. *Observers can decide if they want to consider this a part of begging or not.<br>5. Chicks can beg while sitting down or standing up. | - Begging behavior begins when any of the actions described in the 'Behavior Definition' occur.<br>– **Begging behavior ends when there is an absence ($\leq 3$ seconds) of the actions as described in 'Behavior Definition.'** |
| Aggression | State | 1. Most often carried out by the alpha chick toward the beta chick.<br>* alpha = A chick = first hatched chick within a brood; alpha chicks are marked with red *beta = B chick = second hatched chick within a brood; beta chicks are marked with blue sharpie on top of their head<br>2. Aggressor uses a quick thrusting motion of their head that allows their beak to come into contact with the victim's face area, top of head, or nape. This motion occurs over and over to create a 'pecking' motion.<br>3. Multiple pecks to the face area, top of head and/or nape of victim.<br>4. Pecks can be accompanied by grabbing motion: aggressor will grab<br>skin or feathers in tip of beak and use small, quick, and side-to-side jerking motions (“wringing”). | - Aggression begins when any of the actions described in 'Behavior Definition' occur.<br>– **Aggression ends when there is an absence ($\leq 5$ seconds) of the action as described in 'Behavior Definition'.** |
| 1 Feeding* | Point | *When food **is** visible -*<br>1. Parent opens mouth wide.<br>2. The focal chick inserts its beak into the parent’s mouth, just at the beginning of the parent’s throat.<br>3. Can see chick pull out bits or strings of food and eat them (opens and closes mouth or can see a swallow).<br>OR<br>1. The parent holds a chunk at the tip of its beak.<br>2. The focal chick pecks at the chunk(s) of food. If there is a chewing motion from the chick it is considered a successful feed. Chick may repeat this a few times in a matter of minutes.<br>*When food **is not** visible -*<br>1. Parent opens mouth wide.<br>2. The focal chick inserts its beak far back into the parent’s mouth, just at the beginning of the throat.<br>3. The focal chick pulls the beak out, no food or strings of food are visible, but the chick moves its beak in an eating/chewing motion or swallowing is visible. | - Feeding occurs when any of the actions described in the ‘Behavior Definition’ can be observed.<br>- At least 10 seconds must pass between successful feeding bouts in order to be considered separate feeds h. |
| Unsuccessful Feeding* | Point | 1. If there is an attempt of the parent to feed the focal chick (regurgitation motion from parent or you see food come to the forefront of parent’s bill) but the chick does not take the food or is disrupted when going to take the food, this is considered an unsuccessful feed. | - Unsuccessful feeding occurs when any of the actions described in the ‘Behavior Definition’ can be observed.<br>- At least 10 |
|  |  | OR<br>2. If there is an attempt of the parent to feed the focal chick (regurgitation motion from parent or you see food come to the forefront of parent's bill) and the chick appears to take some of the food but there is NO chewing/swallowing motion from the focal chick then it is considered an unsuccessful feed<br>OR<br>3. The focal chick receives a visible chunk of food from the parent's beak, attempts to swallow it, but the chunk comes back out of the chick's mouth and falls into the nest.<br>OR<br>4. The focal chick receives a visible chunk of food from the parent's beak, but before it can eat off the chunk of food the parent re-swallows the food. | seconds must pass between unsuccessful feeding bouts in order to be considered separate behavior occurrences. Within those 10 or more seconds, any other defined behaviors (including inactivity) may occur and should be coded as such. |

\*Successful and unsuccessful feedings may be combined into a single 'feeding attempt' behavior if the differentiation between the outcome of feeding attempts proves challenging.

## Appendix 2: Model selection tables for Experiment 1 (endogenous corticosterone elevation) that were not included in the main manuscript

Here we report the results for model selection on the analysis done with feeding treatment and its interactions included as predictors. Models with deltaAICc <2 are indicated in bold; models with deltaAICc <2 but greater than the null model are in bold italics. In the tables below we report all models included within a cumulative model weight of 0.95.

**Table A2.1.** Model selection results for **baseline corticosterone** as a function of feeding treatment, behavior, and sex.

| Model | K | AICc | Delta AICc | AICc Wt | Cum .Wt | LL |
| --- | --- | --- | --- | --- | --- | --- |
| **Feeding treatment** | **3** | **27.96** | **0** | **0.37** | **0.37** | **-10.57** |
| Sex+Feeding treatment | 4 | 30.46 | 2.5 | 0.11 | 0.48 | -10.52 |
| Feeding treatment + Begging previous hour | 4 | 30.51 | 2.55 | 0.1 | 0.58 | -10.54 |
| Feeding treatment + Aggression previous hour | 4 | 30.53 | 2.57 | 0.1 | 0.69 | -10.55 |
| Feeding treatment + Feeding previous hour | 4 | 30.56 | 2.59 | 0.1 | 0.79 | -10.56 |
| Sex * Feeding treatment | 5 | 33.12 | 5.16 | 0.03 | 0.82 | -10.45 |
| Feeding treatment * Begging previous hour | 5 | 33.14 | 5.18 | 0.03 | 0.85 | -10.46 |
| Sex + Feeding treatment + Begging previous hour | 5 | 33.21 | 5.24 | 0.03 | 0.87 | -10.49 |
| Sex + Feeding treatment + Aggression previous hour | 5 | 33.22 | 5.25 | 0.03 | 0.9 | -10.5 |
| Sex + Feeding treatment + Feeding previous hour | 5 | 33.26 | 5.29 | 0.03 | 0.93 | -10.52 |
| Feeding treatment * Aggression previous hour | 5 | 33.28 | 5.31 | 0.03 | 0.95 | -10.53 |

**Table A2.2.**
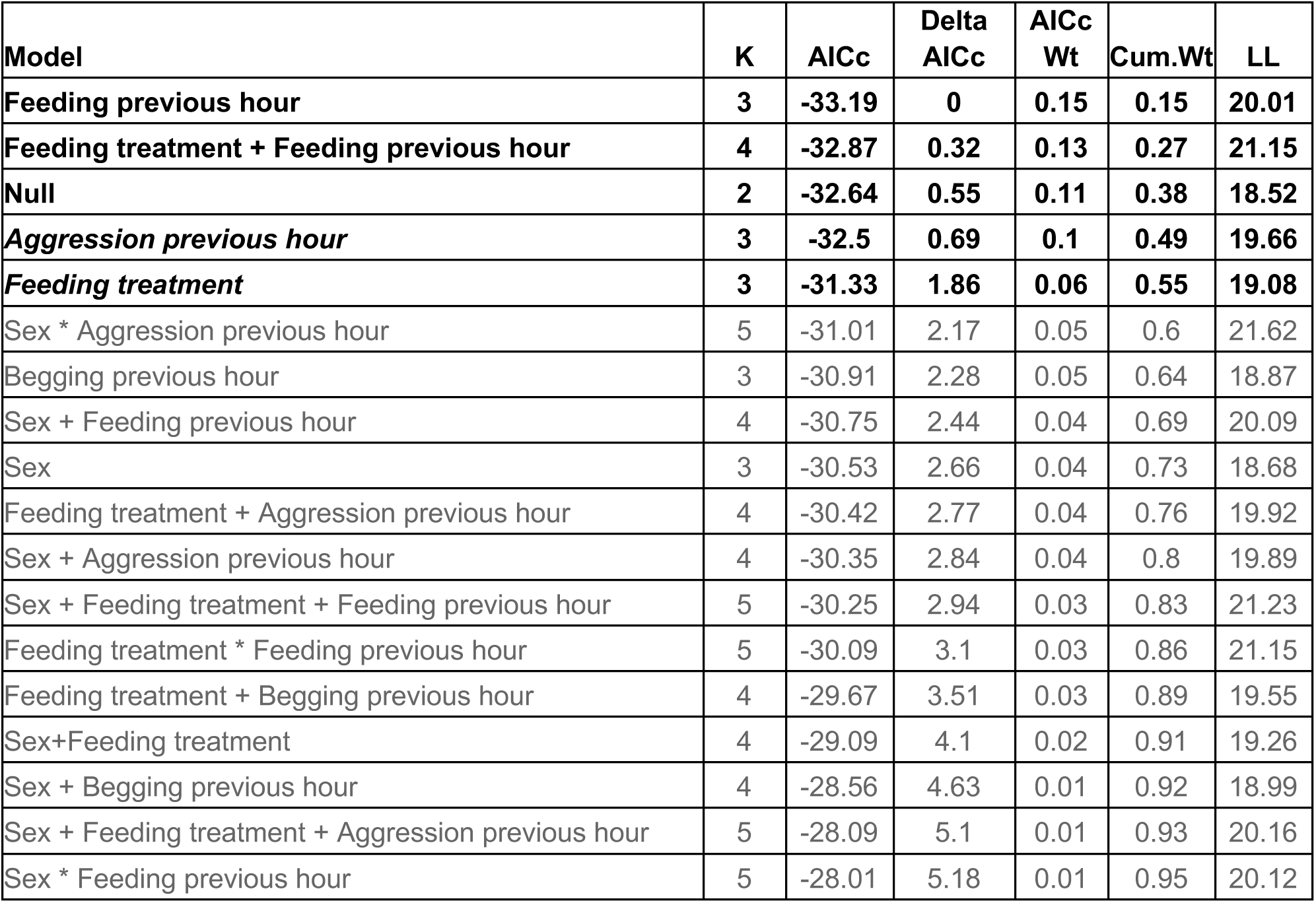
Model selection results for **post-restraint corticosterone** as a function of feeding treatment, behavior, and sex.

| **Model** | **K** | **AICc** | **Delta**<br>**AICc** | **AICc**<br>**Wt** | **Cum.Wt** | **LL** |
| --- | --- | --- | --- | --- | --- | --- |
| **Feeding previous hour** | **3** | **-33.19** | **0** | **0.15** | **0.15** | **20.01** |
| **Feeding treatment + Feeding previous hour** | **4** | **-32.87** | **0.32** | **0.13** | **0.27** | **21.15** |
| **Null** | **2** | **-32.64** | **0.55** | **0.11** | **0.38** | **18.52** |
| ***Aggression previous hour*** | **3** | **-32.5** | **0.69** | **0.1** | **0.49** | **19.66** |
| ***Feeding treatment*** | **3** | **-31.33** | **1.86** | **0.06** | **0.55** | **19.08** |
| Sex * Aggression previous hour | 5 | -31.01 | 2.17 | 0.05 | 0.6 | 21.62 |
| Begging previous hour | 3 | -30.91 | 2.28 | 0.05 | 0.64 | 18.87 |
| Sex + Feeding previous hour | 4 | -30.75 | 2.44 | 0.04 | 0.69 | 20.09 |
| Sex | 3 | -30.53 | 2.66 | 0.04 | 0.73 | 18.68 |
| Feeding treatment + Aggression previous hour | 4 | -30.42 | 2.77 | 0.04 | 0.76 | 19.92 |
| Sex + Aggression previous hour | 4 | -30.35 | 2.84 | 0.04 | 0.8 | 19.89 |
| Sex + Feeding treatment + Feeding previous hour | 5 | -30.25 | 2.94 | 0.03 | 0.83 | 21.23 |
| Feeding treatment * Feeding previous hour | 5 | -30.09 | 3.1 | 0.03 | 0.86 | 21.15 |
| Feeding treatment + Begging previous hour | 4 | -29.67 | 3.51 | 0.03 | 0.89 | 19.55 |
| Sex+Feeding treatment | 4 | -29.09 | 4.1 | 0.02 | 0.91 | 19.26 |
| Sex + Begging previous hour | 4 | -28.56 | 4.63 | 0.01 | 0.92 | 18.99 |
| Sex + Feeding treatment + Aggression previous hour | 5 | -28.09 | 5.1 | 0.01 | 0.93 | 20.16 |
| Sex * Feeding previous hour | 5 | -28.01 | 5.18 | 0.01 | 0.95 | 20.12 |

**Table A2.3.** Model selection results for **change in aggression** as a function of feeding treatment, behavior, sex, and corticosterone.

| Model | K | AICc | Delta AICc | AICc Wt | Cum.Wt | LL |
| --- | --- | --- | --- | --- | --- | --- |
| **Sex** | **3** | **-334.96** | **0** | **0.25** | **0.25** | **170.89** |
| **Log Post-Restraint corticosterone + Sex** | **4** | **-333.21** | **1.74** | **0.1** | **0.35** | **171.32** |
| Log Baseline corticosterone * Sex | 5 | -332.87 | 2.09 | 0.09 | 0.44 | 172.55 |
| Sex + Feeding Treatment | 4 | -332.65 | 2.3 | 0.08 | 0.52 | 171.04 |
| Log Baseline corticosterone + Sex | 4 | -332.37 | 2.59 | 0.07 | 0.59 | 170.9 |
| Sex * Feeding Treatment | 5 | -332.18 | 2.77 | 0.06 | 0.65 | 172.2 |
| Null | 2 | -332.1 | 2.85 | 0.06 | 0.71 | 168.25 |
| Log Post-Restraint corticosterone * Sex | 5 | -331.48 | 3.48 | 0.04 | 0.76 | 171.85 |
| Log Post-Restraint corticosterone | 3 | -330.84 | 4.12 | 0.03 | 0.79 | 168.83 |
| Sex + Feeding Treatment + Log Post-Restraint corticosterone | 5 | -330.56 | 4.39 | 0.03 | 0.82 | 171.39 |
| Sex + Log Baseline corticosterone + Log Post-Restraint corticosterone | 5 | -330.45 | 4.51 | 0.03 | 0.84 | 171.34 |
| Sex + Feeding Treatment + Log Baseline corticosterone | 5 | -330.07 | 4.89 | 0.02 | 0.86 | 171.14 |
| Log Post-Restraint corticosterone * Feeding Treatment | 5 | -329.97 | 4.99 | 0.02 | 0.88 | 171.1 |
| Feeding Treatment | 3 | -329.88 | 5.08 | 0.02 | 0.9 | 168.35 |
| Log Baseline corticosterone | 3 | -329.71 | 5.24 | 0.02 | 0.92 | 168.27 |
| Sex + Feeding Treatment + Log Baseline corticosterone + Sex * Feeding Treatment | 6 | -329.65 | 5.31 | 0.02 | 0.94 | 172.44 |
| Sex + Feeding Treatment + Log Post-Restraint corticosterone + Sex * Feeding Treatment | 6 | -329.49 | 5.47 | 0.02 | 0.96 | 172.36 |

**Table A2.4.** Model selection results for **change in begging** as a function of feeding treatment, behavior, sex, and corticosterone.

| Model | K | AICc | Delta AICc | AICcWt | Cum.Wt | LL |
| --- | --- | --- | --- | --- | --- | --- |
| **Log Post-Restraint corticosterone** | **3** | **-316.97** | **0** | **0.18** | **0.18** | **161.9** |
| **Log Post-Restraint corticosterone + Sex** | **4** | **-316.08** | **0.88** | **0.11** | **0.29** | **162.75** |
| **Null** | **2** | **-315.9** | **1.06** | **0.1** | **0.39** | **160.15** |
| **Sex** | **3** | **-315.51** | **1.45** | **0.08** | **0.48** | **161.17** |
| **Log Baseline corticosterone +Log Post-Restraint corticosterone** | **4** | **-315** | **1.96** | **0.07** | **0.54** | **162.22** |
| Log Post-Restraint corticosterone + Feeding Treatment | 4 | -314.65 | 2.32 | 0.06 | 0.6 | 162.04 |
| Log Post-Restraint corticosterone * Feeding Treatment | 5 | -314.41 | 2.55 | 0.05 | 0.64 | 163.32 |
| Feeding Treatment | 3 | -314.19 | 2.78 | 0.04 | 0.69 | 160.51 |
| Log Post-Restraint corticosterone * Sex | 5 | -314.03 | 2.94 | 0.04 | 0.73 | 163.12 |
| Log Baseline corticosterone | 3 | -313.91 | 3.05 | 0.04 | 0.77 | 160.37 |
| Sex + Log Baseline corticosterone + Log Post-Restraint corticosterone | 5 | -313.87 | 3.1 | 0.04 | 0.8 | 163.04 |
| Sex + Feeding Treatment | 4 | -313.74 | 3.22 | 0.03 | 0.84 | 161.59 |
| Sex + Feeding Treatment + Log Post-Restraint corticosterone | 5 | -313.66 | 3.31 | 0.03 | 0.87 | 162.94 |
| Log Baseline corticosterone * Sex | 5 | -313.44 | 3.53 | 0.03 | 0.9 | 162.83 |
| Log Baseline corticosterone + Sex | 4 | -313.3 | 3.66 | 0.03 | 0.93 | 161.37 |
| Feeding Treatment + Log Baseline corticosterone + Log Post-Restraint corticosterone | 5 | -312.23 | 4.74 | 0.02 | 0.95 | 162.23 |

**Table A2.5.** Model selection results for **change in feeding** as a function of feeding treatment, behavior, sex, and corticosterone.

| Model | K | AICc | Delta AICc | AICc Wt | Cum.Wt | LL |
| --- | --- | --- | --- | --- | --- | --- |
| **Sex** | **3** | **-370.22** | **0** | **0.25** | **0.25** | **188.52** |
| **Log Post-Restraint corticosterone + Sex** | **4** | **-369.92** | **0.29** | **0.22** | **0.47** | **189.68** |
| **Log Post-Restraint corticosterone \* Sex** | **5** | **-369.28** | **0.94** | **0.16** | **0.62** | **190.75** |
| Log Baseline corticosterone + Sex | 4 | -367.64 | 2.58 | 0.07 | 0.69 | 188.53 |
| Sex + Feeding Treatment | 4 | -367.62 | 2.6 | 0.07 | 0.76 | 188.52 |
| Sex + Feeding Treatment + Log Post-Restraint corticosterone | 5 | -367.25 | 2.97 | 0.06 | 0.82 | 189.74 |
| Sex + Log Baseline corticosterone + Log Post-Restraint corticosterone | 5 | -367.14 | 3.08 | 0.05 | 0.87 | 189.68 |
| Sex * Feeding Treatment | 5 | -364.89 | 5.33 | 0.02 | 0.89 | 188.55 |
| Sex + Feeding Treatment + Log Baseline corticosterone | 5 | -364.86 | 5.35 | 0.02 | 0.91 | 188.54 |
| Log Baseline corticosterone * Sex | 5 | -364.85 | 5.37 | 0.02 | 0.92 | 188.54 |
| Log Post-Restraint corticosterone | 3 | -364.34 | 5.87 | 0.01 | 0.94 | 185.58 |
| Sex + Feeding Treatment + Log Baseline corticosterone + Log Post-Restraint corticosterone | 6 | -364.34 | 5.88 | 0.01 | 0.95 | 189.78 |

**Table A2.6.** Model selection results for **B chick days until death (survival duration)** as a function of feeding treatment, sex, and A chick corticosterone.

| Model | K | Delta AICc | AICc Wt | LL | adj R squared |
| --- | --- | --- | --- | --- | --- |
| **A chick post-restraint corticosterone x feeding treatment** | **5** | **0** | **0.45** | **-74.18** | **0.2667** |
| A chick post-restraint corticosterone | 3 | 2.23 | 0.15 | -78.5 | 0.0225 |
| A chick post-restraint corticosterone + sex | 4 | 3.04 | 0.1 | -77.4 | 0.0690 |
| A chick baseline corticosterone | 3 | 3.35 | 0.08 | -79.06 | -0.0287 |
| A chick post-restraint corticosterone * feeding treatment + sex | 6 | 3.54 | 0.08 | -74.02 | 0.2345 |
| A chick baseline corticosterone + sex | 4 | 4.41 | 0.05 | -78.08 | 0.0095 |
| A chick post-restraint corticosterone + feeding treatment | 4 | 5.23 | 0.03 | -78.49 | -0.0284 |
| A chick baseline + feeding treatment | 4 | 5.43 | 0.03 | -78.59 | -0.0374 |

